# A pedigree-based map of crossovers and non-crossovers in aye-ayes (*Daubentonia madagascariensis*)

**DOI:** 10.1101/2024.11.08.622675

**Authors:** Cyril J. Versoza, Audald Lloret-Villas, Jeffrey D. Jensen, Susanne P. Pfeifer

## Abstract

Gaining a better understanding of rates and patterns of meiotic recombination is crucial for improving evolutionary genomic modelling, with applications ranging from demographic to selective inference. Although previous research has provided important insights into the landscape of crossovers in humans and other haplorrhines, our understanding of both the considerably more common outcome of recombination (i.e., non-crossovers) as well as the landscapes in more distantly-related primates (i.e., strepsirrhines) remains limited owing to difficulties associated with both the identification of non-crossover tracts as well as species sampling. Thus, in order to elucidate recombination patterns in this under-studied branch of the primate clade, we here characterize crossover and non-crossover landscapes in aye-ayes utilizing whole-genome sequencing data from six three-generation pedigrees as well as three two-generation multi-sibling families, and in so doing provide novel insights into this important evolutionary process shaping genomic diversity in one of the world’s most critically endangered primate species.

## INTRODUCTION

Recombination is a fundamental biological process required for faithful gametogenesis in most sexually reproducing species (see the reviews of Baudat et al. 2013; Johnston 2024). Apart from being essential for the proper pairing of homologous chromosomes and their segregation into gametes during meiosis (Keeney 2001), recombination also plays an important evolutionary role in shuffling genetic variation, improving the efficacy of natural selection by breaking linkage interference between segregating beneficial and deleterious alleles (Hill and Robertson 1966; Felsenstein 1974; and see the review of Charlesworth and Jensen 2021).

In primates, as in many other organisms, meiotic recombination predominantly occurs in 1-2 kb long regions of the genome – so called recombination “hotspots” – the location of which is mainly determined by the zinc-finger protein PRDM9 (Baudat et al. 2010; Myers et al. 2010; Parvanov et al. 2010). By binding specific DNA sequence motifs and trimethylating histone H3 at lysines 4 and 36 (Powers et al. 2016), PRDM9 guides the meiotic machinery to initiate the formation of DNA double-strand breaks, the repair of which may result in either a reciprocal exchange between homologs (termed a crossover; CO) or a unidirectional replacement of a genomic region in one chromosome leaving the donor homolog unmodified (termed a non-crossover; NCO) (see reviews of Wang et al. 2015; Lorenz and Mpaulo 2022; Johnston 2024).

While both COs and NCOs play important roles in shaping genetic diversity (Przeworski and Wall 2001), previous studies have suggested that NCOs tend to be considerably more common than COs in most organisms (Jeffreys and May 2004; Cole et al. 2010; Comeron et al. 2012; Li et al. 2019). However, owing to the difficulty in detecting the often very small NCO events (with most tract lengths in humans being < 1 kb; Jeffreys and May 2004; Williams et al. 2015; Halldorsson et al. 2016), combined with the need for a segregating variant to be present in the donor homolog in order to allow identification (i.e., the tract remains undetectable if the donor and converted sequence are identical), the NCO landscape remains comparatively understudied. For these same reasons, combined with other underlying assumptions such as geometrically-distributed tract lengths, computational approaches for NCO inference are generally characterized by poor accuracy (see discussion of Wall et al. 2022). As such, time-consuming and costly direct pedigree-sequencing studies remain the most promising avenue for improving our understanding of this process (and see Peñalba and Wolf 2020 for a detailed overview of current methodologies).

With regards to primates specifically, although previous research has begun to elucidate the rates and patters of recombination in haplorrhines – a suborder of primates that includes humans (Kong et al. 2002, 2010; Coop et al. 2008; Pratto et al. 2014; Williams et al. 2015; Halldorsson et al. 2016), other great apes (Auton, Fledel-Alon, Pfeifer, Venn et al. 2012; Pfeifer and Jensen 2016; Stevison et al. 2016), and monkeys (Rogers et al. 2000, 2006; Cox et al. 2006; Jasinska et al. 2007; Xue et al. 2016, 2020; Pfeifer 2020; Wall et al. 2022; Versoza, Weiss et al. 2024) – little remains known about this process in strepsirrhines. As the most basal suborder of primates, gaining insights into the population genetic forces shaping the genomes of strepsirrhines is crucially important, not only to improve our understanding of primate evolution, but also for elucidating the scale at which recombination rates and spatial distributions of CO and NCO events may change between taxonomic groups, as this variation has been shown to be substantial (see reviews of Paigen and Petkov 2010; Stapley et al. 2017). Given the vital role of recombination in maintaining genetic diversity, such inference is additionally important for the development of effective conservation strategies, particularly as many strepsirrhines are highly endangered (Gross 2017). For example, amongst the more than 100 species of strepsirrhines endemic to Madagascar (Mittermeier et al. 2010), aye-ayes (*Daubentonia madagascariensis*) are one of the most threatened by anthropogenic activities, such as slash-and-burn agriculture, logging, mining, and urbanization (Suzzi-Simmons 2023), with the resulting extensive habitat loss and fragmentation having already decimated their populations to an estimated 1,000 to 10,000 individuals (Louis et al. 2020).

Thus, to elucidate the rates and patterns of recombination in this under-studied branch of the primate tree – which represents an early split in the primate clade and thus a valuable comparative outgroup for future primate studies – we here investigate both the more commonly studied CO and the less-commonly studied NCO landscapes in aye-ayes. Using whole-genome sequencing data from six three-generation pedigrees and three two-generation multi-sibling families, we present the first recombination rate estimates for a strepsirrhine and thereby provide novel insights into this important evolutionary process shaping genetic diversity in one of the world’s most critically endangered primate species.

## RESULTS AND DISCUSSION

To study the rates and patterns of recombination in aye-ayes, 14 individuals were selected from a multi-generation pedigree housed at the Duke Lemur Center, the genomes of which were sequenced to mean depths of 50X (Supplementary Table 1; Versoza et al. 2024a). After mapping reads to the species-specific genome assembly (Versoza and Pfeifer 2024), autosomal variants were called following the Genome Analysis Toolkit’s Best Practices (van der Auwera and O’Connor 2020), and filtered using a set of coverage-, genotype-, and inheritance-based criteria described in Versoza, Weiss et al. (2024), resulting in a high-confidence call set consisting of 1.8 million variants (Supplementary Table 2). This call set was divided into six three-generation pedigrees and three two-generation nuclear families with multiple offspring (Supplementary Figure 1) for which gamete transmission could be tracked in order to identify recombination events based on “phase-informative” markers – that is, heterozygous variants for which the parent-of-origin could be determined (with an average of 0.5 million phase-informative markers per pedigree / family; Supplementary Table 3, and see Figure 1b in Versoza, Weiss et al. 2024 and Supplementary Figure 2 for a schematic of the workflow in the three-generation pedigrees and two-generation nuclear families; respectively). Based on these phase-informative markers, recombination events were classified as either COs – that is, a single change of haplotype phase along a chromosome (from maternally-inherited to paternally-inherited haplotype blocks or *vice versa*) – or NCOs – that is, phase-informative markers that mismatched surrounding haplotype blocks (for additional details, see “Materials and Methods” in the Supplementary Materials).

**Figure 1.**
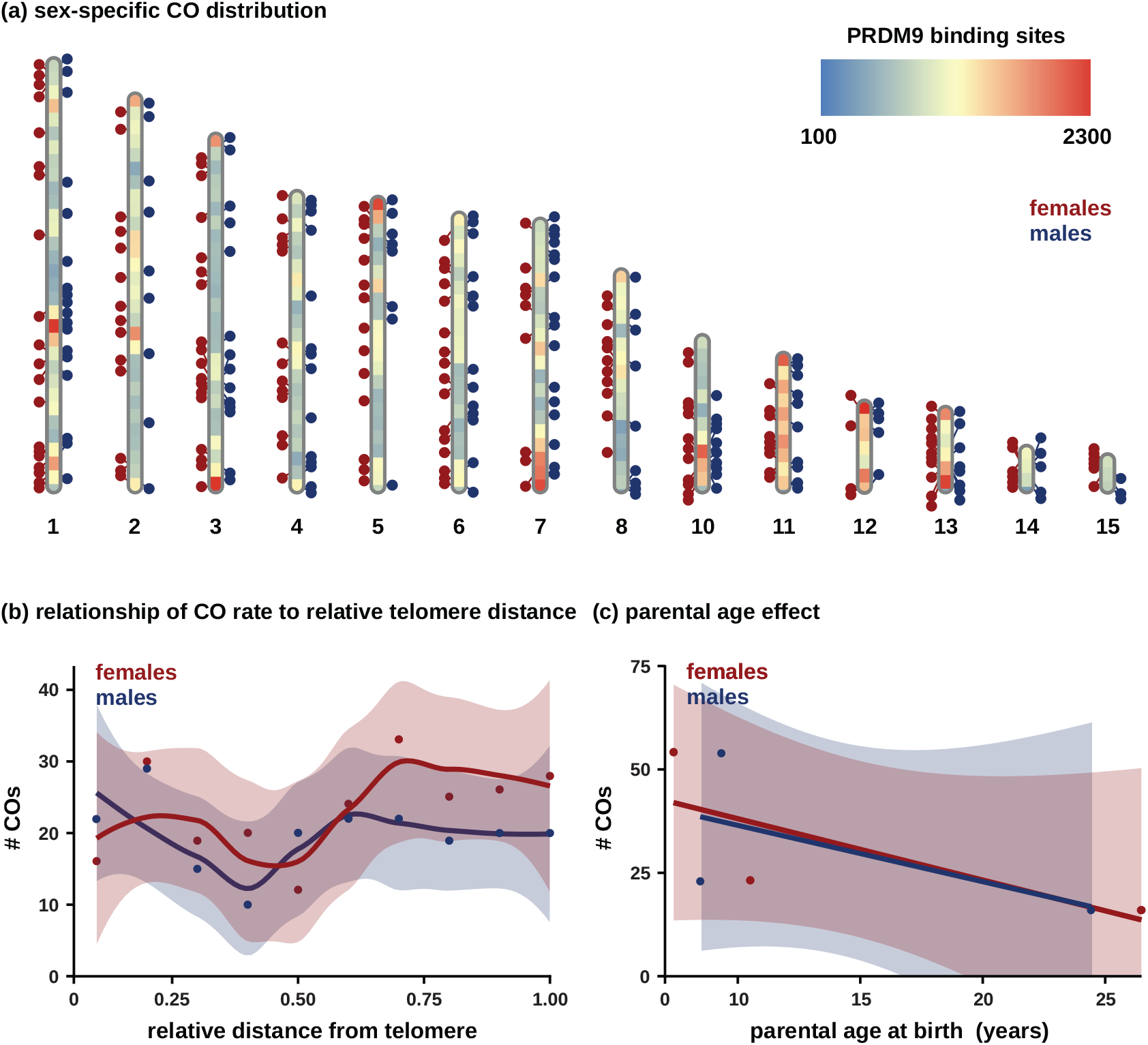
Characteristics of crossovers. (a) The genomic distribution of CO events in females (shown as red circles) and males (blue circles) across the autosomes (note that scaffold 9, i.e., chromosome X, is not displayed). The heatmap indicates the density of PRDM9 binding sites across 10 Mb genomic regions. (b) Relationship between female and male CO rates and relative proximity to telomeric regions. (c) Relationship between maternal and paternal age at birth and the number of CO events.

### The landscape of COs in aye-ayes

A total of 117 and 310 putative CO events were identified across the autosomes in 6 and 18 meioses through the pedigree-based and family-based approaches, respectively. Visual inspection of these initial datasets revealed a clustering (i.e., ≥ 2 COs originating from the same meiosis located within < 1 Mb) of 36 COs across 8 genomic regions, with the majority of these 36 events (94.4%) identified through only one of the two approaches (Supplementary Table 4). Although COs form at random during meiosis, crossover interference tightly regulates their location within each chromosome (Muller 1916; Broman and Weber 2000), making such a pattern highly implausible in nature (see the discussions of Wall et al. 2022; Versoza, Weiss et al. 2024). More likely, this observation is driven by genotyping errors; thus, these CO events were excluded from further analyses, as is common practice. Subsequently, pedigree-based and family-based datasets were consolidated across the 20 meioses, and an additional 17 COs were removed either because the event was detected across multiple meioses (16 COs), possibly indicating a mis-placement or inversion of a contig during the genome assembly, or because it occurred at the same position as a NCO in another individual (1 CO), thus likely resulting from a genotyping error. The final dataset contained 305 COs: 163 and 142 COs in the 10 maternal and 10 paternal meioses, respectively (Figure 1a; and see Supplementary Table 5 for details).

Aye-ayes exhibit one of the lowest levels of nucleotide diversity of any primate studied to date (Perry et al. 2013; Terbot et al. 2024; Soni et al. 2024) – yet, despite the resulting lower marker density across the pedigree, the median resolution of male and female CO events (12.9 and 20.0 kb; Supplementary Figure 3, and see Supplementary Table 6 for a summary) is on par with those previously obtained from similarly-sized non-human primate pedigree studies (7.7 and 22.3 kb in olive baboons [Wall et al. 2022] and rhesus macaques [Versoza, Weiss et al. 2024], respectively). In accordance with crossover assurance, ensuring an obligate CO between homologous chromosomes (or chromosome arms) during meiosis (Jones and Franklin 2006), an average of 1.1 COs were identified per chromosome and meiosis. In total, between 10 and 21 COs were observed per meiosis across the 14 autosomes (Supplementary Table 6), with CO density in males and females roughly inversely correlated with chromosome size (Supplementary Figure 4). CO events were significantly enriched in regions harboring predicted PRDM9 binding sites (210 out of 305 COs, or 68.9%; *p*-value < 2.2e-16, one-sided binomial test; Figure 1a) using the degenerate sequence motifs previously determined in great apes (Berg et al. 2011; Auton, Fledel-Alon, Pfeifer, Venn et al. 2012; Schwartz et al. 2014; Stevison et al. 2016) as a proxy (note that given the high turn-over rate of PRDM9 binding motifs between species, the actual overlap is likely even greater). Consistent with a preferential binding of PRDM9 in intronic and intergenic regions that are often more accessible in chromatin structure (Coop et al. 2008; Walker et al. 2015), CO events were enriched in these genomic regions (Supplementary Figure 5). Additionally, a clustering of COs was observed toward the telomeric ends in males, while females displayed an overall larger number of COs that were more evenly spaced throughout the genome (Figure 1b), as previously observed in other primates (e.g., Kong et al. 2002, 2010; Coop et al. 2008; Wall et al. 2022; and see Lenormand and Dutheil 2005 and Sardell and Kirkpatrick 2020 for a discussion of this widespread phenomenon). Moreover, in agreement with recent studies in humans (Porubsky et al. 2024) and chimpanzees (Venn et al. 2014), the frequency of CO events decreased with both maternal and paternal age in aye-ayes (with 1.48 and 1.36 fewer COs per year, respectively; Figure 1c); however, given the small sample size, this observation is not statistically significant (maternal: adjusted *R*^*2*^ = 0.1307, *p*-value: 0.4583 and paternal: adjusted *R*^*2*^ = -0.2682, *p*-value: 0.5864). Based on the number of COs per meiosis, the sex-averaged autosomal genetic map was estimated to be 1,525 cM in length (Table 1) – approximately 25-35% shorter than those of catarrhines (Rogers et al. 2000, 2006; Cox et al. 2006; Jasinska et al. 2007; Wall et al. 2022; Versoza, Weiss et al. 2024) as may be anticipated from the lower karyotype (2*n* = 30 in aye-ayes [Tattersall 1982] vs. 2*n* = 42 and 60 in rhesus macaques [Owen et al. 2016] and vervet monkeys [Finelli et al. 1999], respectively) (Pardo-Manuel de Villena and Sapienza 2001), which, in turn, exhibit shorter map lengths than hominoids (Kong et al. 2002, 2010; Venn et al. 2014). Similar to other primates, females exhibit an overall longer genetic map length than males (1,630 cM vs 1,420 cM) – however, the ratio of the female to male autosomal map length (1.15) is lower than that observed in humans (1.36; Porubsky et al. 2024). The genome-wide average CO rates in males and females were thus estimated to be 0.77 cM/Mb and 0.94 cM/Mb (Table 1) – approximately 40-50% lower than the average rates of 1.3 cM/Mb and 2.0 cM/Mb reported in humans (Bhérer et al. 2017) – with the overall lower sex-averaged rate (0.85 cM/Mb) likely contributing to the low levels of genetic diversity observed in the species (Perry et al. 2013; Terbot et al. 2024; Soni et al. 2024).

**Table 1.** Number of crossover (CO) events and genetic distances identified in maternal / paternal meioses.

| **scaffold** | **length (Mb)** | **# CO events** | **cM** | **cM/Mb** |
| --- | --- | --- | --- | --- |
| 1 | 316.17 | 20 / 18 | 200 / 180 | 0.63 / 0.57 |
| 2 | 290.59 | 14 / 9 | 140 / 90 | 0.48 / 0.31 |
| 3 | 261.42 | 18 / 14 | 180 / 140 | 0.69 / 0.54 |
| 4 | 219.69 | 13 / 14 | 130 / 140 | 0.59 / 0.64 |
| 5 | 215.45 | 14 / 8 | 140 / 80 | 0.65 / 0.37 |
| 6 | 204.02 | 16 / 13 | 160 / 130 | 0.78 / 0.64 |
| 7 | 199.60 | 9 / 16 | 90 / 160 | 0.45 / 0.80 |
| 8 | 162.77 | 11 / 9 | 110 / 90 | 0.68 / 0.55 |
| 10 | 114.90 | 12 / 10 | 120 / 100 | 1.04 / 0.87 |
| 11 | 102.08 | 9 / 10 | 90 / 100 | 0.88 / 0.98 |
| 12 | 67.30 | 4 / 5 | 40 / 50 | 0.59 / 0.74 |
| 13 | 62.73 | 11 / 8 | 110 / 80 | 1.75 / 1.28 |
| 14 | 34.25 | 6 / 5 | 60 / 50 | 1.75 / 1.46 |
| 15 | 28.26 | 6 / 3 | 60 / 30 | 2.12 / 1.06 |
| **autosomal**<br>**(sex-averaged)** | **2,279.23** | **163 / 142**<br>**(152.5)** | **1,630 / 1,420**<br>**(1,525)** | **0.94 / 0.77**<br>**(0.85)** |

### The landscape of NCOs in aye-ayes

A total of 151 and 198 putative NCO events were identified through the pedigree-based and family-based approaches, respectively (note that complex events involving multiple, non-contiguous NCO tracts within < 5 kb as well as NCOs with tracts > 10 kb were excluded from analyses as such events were previously shown to frequently represent incorrect genotype calls or assembly errors; Smeds et al. 2016; Wall et al. 2022). After merging the datasets from the pedigree-based and family-based approaches, 88 NCOs were removed, either because the event was detected across multiple meioses (86 NCOs) or because the phase-informative markers overlapped with a structural variant (1 NCO; Versoza et al. 2024b) or CO in another individual (1 NCO). Thus, the final dataset contained 200 NCOs, 95 and 105 of maternal and paternal origin, respectively (Figure 2a; and see Supplementary Table 7 for details).

**Figure 2.**
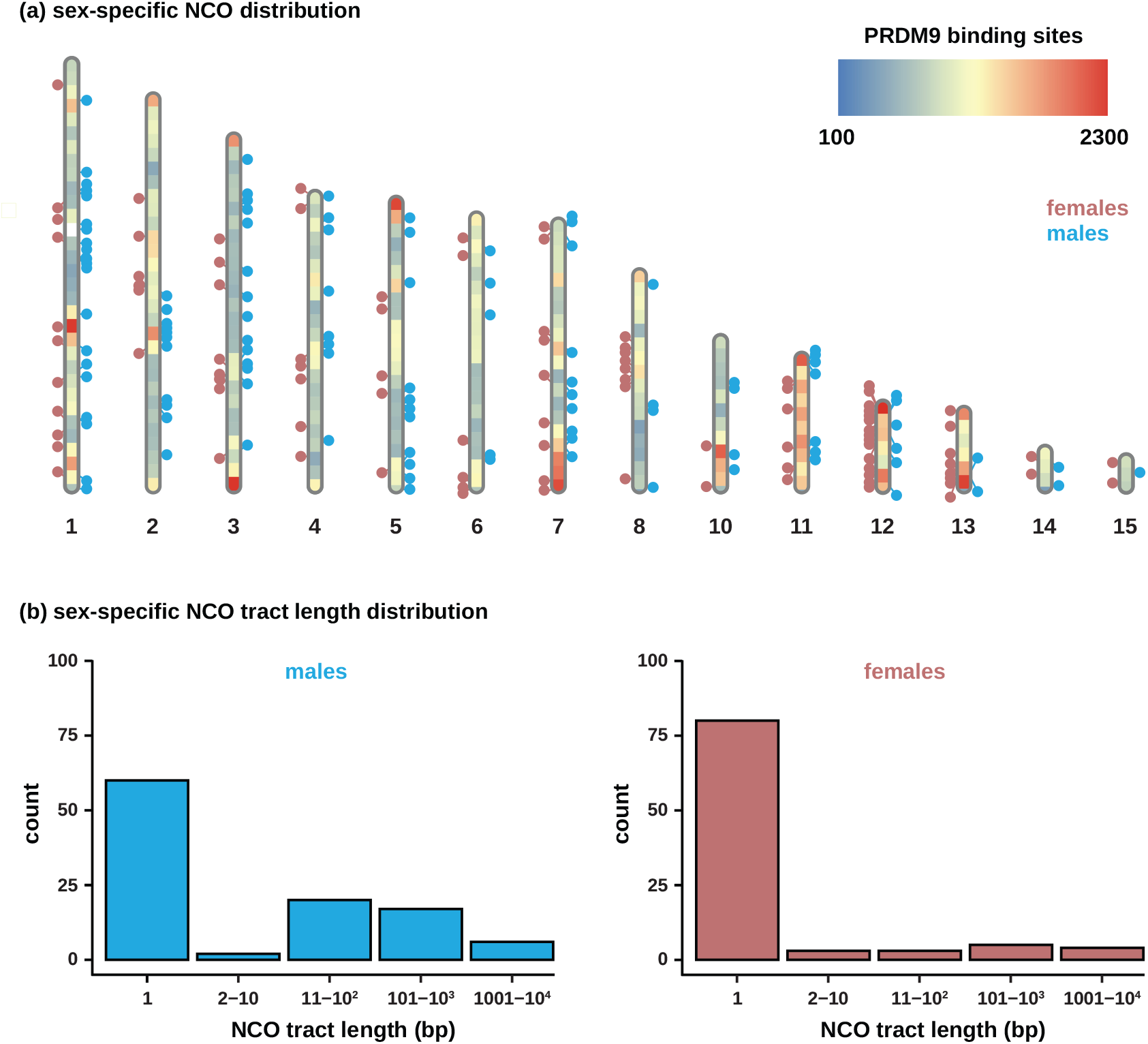
Characteristics of non-crossovers. (a) The genomic distribution of NCO events in females (shown as pink circles) and males (blue circles) across the autosomes (note that scaffold 9, i.e., chromosome X, is not displayed). The heatmap indicates the density of PRDM9 binding sites across 10 Mb genomic regions. (b) Distribution of sex-specific NCO tract lengths (reported in base pairs [bp]) based on phase-informative sites that mismatched the surrounding haplotype block.

On average, 0.71 NCOs were identified per chromosome and meiosis, with 1 to 20 NCOs per meiosis across the 14 autosomes (Supplementary Table 6). Although quantitatively similar to recent estimates in catarrhines (e.g., 1.06 NCOs per chromosome and meiosis in rhesus macaques; Versoza, Weiss et al. 2024), this represents a conservative estimate, both because NCO events occurring between phase-informative markers will inevitably be missed (a particular challenge for species exhibiting low heterozygosity – and thus low marker density – such as aye-ayes) but also because the application of stringent quality metrics required to filter out false positives may have inadvertently removed genuine NCO events.

NCOs are slightly less abundant in females than males (8.6 vs. 9.6) and exhibit shorter average tract lengths (157 vs. 165 bp; Figure 2b and see Supplementary Table 6). The sex-averaged mean tract length of 161 bp is similar to those observed in pedigree studies of other primates (with estimated mean tract length of 55-290 bp in humans [Jeffreys and May 2004], 42-167 bp in baboons [Wall et al. 2022], and 155 bp in rhesus macaques [Versoza, Weiss et al. 2024]). In fact, the majority of events (57.1%) include only a single phase-informative marker, exhibiting a tract length of 1 bp. However, similar to humans (Williams et al. 2015), baboons (Wall et al. 2022), and rhesus macaques (Versoza, Weiss et al. 2024) several NCOs with tract length longer than 1 kb were also observed (6 in males and 4 in females; Supplementary Table 7). Furthermore, in agreement with empirical patterns observed in other primates (Williams et al. 2015; Halldorsson et al. 2016; Wall et al. 2022; Versoza, Weiss et al. 2024), the distribution of NCO tract lengths (Figure 2b) appears more consistent with a power-law (or heavy-tailed) distribution than the single geometric distribution frequently modelled (Frisse et al. 2001; Gay et al. 2007; Yin et al. 2009).

In contrast to the transmission bias towards strong (C and G) alleles observed at NCOs in haplorrhines (68% in humans [Odenthal-Hesse et al. 2014; Williams et al. 2015; Halldorsson et al. 2016], 57.6% in baboons [Wall et al. 2022], and 56.5% in rhesus macaques [Versoza, Weiss et al. 2024]), there is no support for GC-biased gene conversion (see review of Duret and Galtier 2009) at NCOs in aye-ayes (45.7%; 95% CI: 37.5-54.0%; *p*-value = 0.1763; two-sided binomial test) – however, the small sample size limits statistical power in these comparisons.

Overall, aye-ayes exhibit a sex-averaged NCO rate of 6.8 × 10^−7^ per base pair per generation (95% CI: 2.9 × 10^−7^ – 1.1 × 10^−6^) – an order of magnitude lower than the rates previously observed in humans (2.6 × 10^−6^ – 5.2 × 10^−5^; Jeffreys and May 2004; Williams et al. 2015; Halldorsson et al. 2016) and baboons (7.52 × 10^−6^; Wall et al. 2022). Based on an overall autosomal genome length of 2.28 Gb, it is thus expected that an average of ∼1.5 kb will be affected by NCOs in each generation.

In summary, the landscape of CO and NCO events in aye-ayes presented here provides novel insights into a parameter crucial for improving population genetic modelling in this highly endangered species, and will serve as a valuable resource for future comparative genomic studies seeking to understand the long-term evolution of recombination landscapes across the primate clade.

## ACKNOWLEDGEMENTS

We would like to thank Erin Ehmke, Kay Welser, and the Duke Lemur Center for providing the aye-aye samples used in this study. DNA extraction, library preparation, and Illumina sequencing were conducted at Azenta Life Sciences (South Plainfield, NJ, USA). Computations were performed on the Sol supercomputer at Arizona State University (Jennewein et al. 2023). This is Duke Lemur Center publication # XXX.

