## Supplementary Materials for "A pedigree-based map of crossovers and non-crossovers in aye-ayes (*Daubentonia madagascariensis*)"

### MATERIALS AND METHODS

#### Sequencing, read mapping, and variant calling

A total of 14 aye-aye individuals (eight females and six males) were selected from six three-generation pedigrees and three two-generation multi-sibling families available at the Duke Lemur Center, the genomes of which were sequenced to an average coverage of 50X per individual (Supplementary Table 1). Sequencing reads were mapped to the aye-aye genome assembly (DMad_hybrid; GenBank accession number: JBFSEQ000000000; Versoza and Pfeifer 2024) in order to call autosomal, biallelic single nucleotide polymorphisms (SNPs) genotyped in all individuals (see Versoza et al. 2024a for details). In the absence of a curated "gold standard" variant dataset that could serve as a basis for variant filtration in the species, initial calls were filtered using the Genome Analysis Toolkit's Best Practice "hard" filter criteria for germline variants (van der Auwera and O'Connor 2020). Following the methodology described by Versoza, Weiss et al. (2024), a set of additional coverage-, genotype-, and inheritance-based filters was applied to further increase the precision of the variant call set. Specifically, variants were excluded if (i) they were located within repetitive and low-complexity regions of the genome prone to base calling errors and mis-mappings (Pfeifer 2017), or if they exhibited (ii) a total depth of coverage (DP) of less than half or more than twice the average autosomal coverage for each sample, (iii) a genotype quality (GQ) less than 30 (corresponding to a probability of a genotyping error of more than $10^{-3}$), (iv) an excess of heterozygosity (defined here as a $p$-value of 0.01), and/or (v) violated the patterns expected by Mendelian inheritance (as determined using the BCFtools +*mendelian2* plugin [Danecek et al. 2021] together with the information of the pedigree). Additionally, to further reduce variant calling and/or genotyping errors that could potentially lead to spurious recombination events, SNPs located within either (i) a cluster of variants (defined here as ≥ 3 SNPs within a 10 bp window), (ii) 10 bp of an insertion/deletion, or (iii) 2 Mb from the chromosome ends, were also excluded from further analyses. The resulting high-confidence call set consisted of 1,830,431 SNPs with a transition-transversion ratio of 2.62 (Supplementary Table 2).

**Identification of crossover and non-crossover events**

The high-confidence call set was divided into six three-generation pedigrees and three two-generation nuclear families with multiple offspring (Supplementary Figure 1) for which gamete transmission could be tracked in order to identify recombination events based on "phase-informative" markers – that is, heterozygous SNPs for which the parent-of-origin could be determined (Supplementary Table 3). In brief, in the three-generation pedigrees, phase-informative markers were sites at which the $P_0$ individuals exhibited non-identical genotypes, their $F_1$ offspring was heterozygous, and either the $F_1$'s partner or their joint $F_2$ offspring was homozygous (for a schematic of the workflow, see Figure 1b in Versoza, Weiss et al. 2024). In the two-generation nuclear families, maternally phase-informative markers exclusively included those for which the dam was heterozygous and the sire homozygous, whereas paternally phase-informative markers included sites at which the sire was heterozygous and the dam homozygous. Following the methodology outlined in Coop et al. (2008), recombination events were then identified by comparing these phase-informative markers across siblings using a randomly assigned offspring as "template" (note that although recombination events can be identified in a single sibling pair using this approach, it is not possible to unequivocally assign the meiosis in which the event occurred; see Supplementary Figure 2 for a schematic of the workflow). In both cases, heterozygous genotype calls in phase-informative markers required the support of more than 25% but less than 75% of mapped reads to limit genotyping errors in the downstream analyses.

Based on these phase-informative markers, recombination events were classified as either COs – that is, a single change of phase along a chromosome (from maternally-inherited to paternally-inherited haplotype blocks or *vice versa*, ignoring single phase-informative markers within a region of consistent phase) – or NCOs – that is, phase-informative markers that mismatched surrounding haplotype blocks. CO interference prevents the occurrence of two CO events in close proximity to each other (Otto and Payseur 2019), thus following Venn et al. 2014, a minimum distance of 1 Mb was required between two COs that originated from the same meiosis (see Supplementary Table 4 for a summary of the removed CO events). Similarly, following Smeds et al. 2016, to reduce the number of spurious NCO events resulting from genotyping errors, a minimum distance of 5 kb was required between two consecutive NCOs; additionally, complex NCO events longer than 10 kb were excluded from further analyses as previous work demonstrated that these frequently represent assembly errors (Wall et al. 2022). Subsequently, recombination events in an individual detected by both the three-generation pedigree and two-generation family approaches were manually consolidated (only CO events called by both approaches were included in the final dataset); in case of a partial overlap (i.e., sharing of either start or end coordinates), the CO events with the shortest resolution and the NCO events with the longest length tract were selected). In brief, all but one CO events could be identified by the three-generation pedigree and two-generation family approaches in the four shared meiosis – 54% of which shared the start and end coordinates of the break point whereas 46% exhibited partially overlapping coordinates owing to different phase-informative markers in the two approaches. In the latter case, shorter CO resolution tracts were generally observed using the pedigree-based approach (only 24% of events exhibited shorter resolutions in the family-based approach) due to a larger number of phase-informative markers. In contrast, only 40% of NCO events were detected with both approaches (83% with identical break point coordinates), further highlighting the challenge of NCO detection. Lastly, to guard against assembly errors, CO and NCO events detected in more than one meiosis were removed.

**Annotation**

High-confidence COs (defined here as COs with a resolution shorter than 5 kb) and NCOs (with a tract length shorter than 5 kb) were classified by genomic region (i.e., intergenic, upstream, exonic, exonic noncoding RNA [ncRNA], intronic, intronic ncRNA, 3' and 5' UTR, and downstream) and repeat content using ANNOVAR v.2020-06-08 (Wang et al. 2010) together with information from the aye-aye genome annotation (Versoza and Pfeifer 2024). CO and NCO events were checked for overlap with structural variants catalogued in the individuals of this study (Versoza et al. 2024b) as well as any PRDM9 motifs previously identified in humans, bonobos, chimpanzees, and gorillas (Berg et al. 2011; Auton, Fledel-Alon, Pfeifer, Venn et al. 2012; Schwartz et al. 2014; Stevison et al. 2016). Statistics were calculated and plotted using R v.4.2.2 (R Core Team 2022) together with the RIdeogram v.0.2.2 package (Hao et al. 2020).

**Supplementary Table 1**. Sample information, including sequencing coverages.

| pedigree ID | sex | date of birth (yyyy-mm-dd) | coverage |
|---|---|---|---|
| 1 | M | 1986-12-19 | 50.5 |
| 2 | F | 1988-11-30 | 50.2 |
| 3 | M | 1985-12-18 | 53.7 |
| 4 | F | 1983-12-04 | 52.5 |
| 5 | M | – | 54.1 |
| 6 | M | – | 53.7 |
| 7 | M | – | 54.1 |
| 8 | F | – | 54.5 |
| 9 | M | – | 51.8 |
| 10 | F | – | 52.7 |
| 11 | F | – | 52.0 |
| 12 | F | – | 48.5 |
| 13 | F | – | 52.4 |
| 14 | F | – | 52.1 |

**Supplementary Table 2**. The high-confidence variant call set consisted of a total of 1,830,431 autosomal, biallelic, single nucleotide polymorphisms (SNPs) with a transition-transversion ratio (Ts/Tv) of 2.62.

| scaffold | length | # SNPs | Ts/Tv |
|---|---|---|---|
| 1 | 316,165,773 | 254,225 | 2.55 |
| 2 | 290,592,686 | 227,280 | 2.57 |
| 3 | 261,424,170 | 216,555 | 2.51 |
| 4 | 219,686,500 | 172,783 | 2.55 |
| 5 | 215,448,047 | 167,698 | 2.59 |
| 6 | 204,016,426 | 164,129 | 2.66 |
| 7 | 199,604,927 | 160,044 | 2.63 |
| 8 | 162,769,830 | 129,061 | 2.59 |
| 10 | 114,896,738 | 88,836 | 2.72 |
| 11 | 102,076,017 | 82,028 | 2.96 |
| 12 | 67,301,774 | 54,750 | 3.00 |
| 13 | 62,733,483 | 54,138 | 2.89 |
| 14 | 34,254,822 | 33,199 | 2.79 |
| 15 | 28,257,198 | 25,705 | 2.77 |
| ∑ or ∅ | **2,279,228,391** | **1,830,431** | **2.62** |

**Supplementary Table 3**. The number of phase-informative markers in the six three-generation pedigrees and three two-generation nuclear families with multiple offspring.

| **three-generation pedigree**<br>*focal individual – offspring* | **# phase-informative markers** | |
|---|---|---|
| 7 – 12 | 566,042 | |
| 7 – 13 | 571,838 | |
| 8 – 12 | 575,898 | |
| 8 – 13 | 582,556 | |
| 8 – 14 | 590,199 | |
| 9 – 14 | 584,470 | |
| **two-generation family**<br>*sire – dam (# offspring)* | **# phase-informative markers** | |
| | **maternally** | **paternally** |
| 1 – 2 (3) | 421,135 | 488,880 |
| 3 – 4 (4) | 403,532 | 534,380 |
| 7 – 8 (2) | 445,960 | 449,098 |

**Supplementary Table 4**. Details of the recombination events removed in the six three-generation pedigrees and three two-generation nuclear families with multiple offspring. Alternating white and gray shading indicates events that were clustered in the same genomic region.

| scaffold | last marker phase A | first marker phase B | last marker phase B | first marker phase A | reason for removal | # markers btw phases A & B | parent – offspring | approach |
|---|---|---|---|---|---|---|---|---|
| 1 | 306,350,133 | 306,355,042 | 306,385,103 | 306,440,213 | • 2 COs < 1Mb & NCO > 10kb<br>• 2 NCOs < 5kb | 2 | 8 – unassigned | family |
| 2 | 75,042,923 | 75,384,570 | 75,516,808 | 76,968,582 | • 2 COs < 1Mb & NCO > 10kb | 0 | 4 – 9 | family |
| 2 | 139,494,657 | 139,509,358 | 139,530,700 | 139,534,486 | • 2 COs < 1Mb & NCO > 10kb | 44 | 8 – 13 | pedigree + family |
| 3 | 158,459,724 | 163,677,209 | 163,907,831 | 166,854,140 | • 2 COs < 1Mb & NCO > 10kb | 0 | 4 – 10 | family |
| 3 | 163,492,972 | 163,778,328 | 163,862,003 | 163,964,064 | • 2 COs < 1Mb & NCO > 10kb | 4 | 8 – 12, 8 – 13 | pedigree |
| 3 | 163,582,977 | 163,773,215 | 163,926,177 | 163,950,022 | • 2 COs < 1Mb & NCO > 10kb | 17 | 1 – 6 | family |
| 3 | 163,615,284 | 163,782,477 | 163,945,287 | 163,955,498 | • 2 COs < 1Mb & NCO > 10kb | 6 | 9 – 14 | pedigree |
| 3 | 163,677,209 | 163,778,328 | 163,917,499 | 163,959,863 | • 2 COs < 1Mb & NCO > 10kb | 2 | 2 – 6 | family |
| 3 | 163,677,209 | 163,782,477 | 163,945,287 | 163,955,498 | • 2 COs < 1Mb & NCO > 10kb | 8 | 7 – 13 | pedigree |
| 3 | 163,757,163 | 163,782,477 | 163,945,287 | 163,955,498 | • 2 COs < 1Mb & NCO > 10kb | 8 | 7 – 12 | pedigree |
| 7 | 6,737,471 | 6,951,451 | 6,965,766 | 6,972,541 | • 2 COs < 1Mb & NCO > 10kb | 0 | 7 – unassigned | family |
| 8 | 55,622,158 | 55,661,940 | 55,683,032 | 55,683,956 | • 2 COs < 1Mb & NCO > 10kb | 27 | 9 – 14 | pedigree |
| 8 | 55,622,158 | 55,664,152 | 55,683,032 | 55,683,956 | • 2 COs < 1Mb & NCO > 10kb | 27 | 7 – 13 | pedigree |
| 8 | 55,622,158 | 55,664,152 | 55,683,050 | 55,683,956 | • 2 COs < 1Mb & NCO > 10kb | 30 | 7 – unassigned | family |
| 8 | 55,658,359 | 55,664,331 | 55,683,050 | 55,707,692 | • 2 COs < 1Mb & NCO > 10kb<br>• 2 NCOs < 5kb | 27 | 1 – 6 | family |
| 11 | 90,020,277 | 90,092,273 | 90,126,449 | 90,148,047 | • 2 COs < 1Mb & NCO > 10kb | 0 | 2 – 5 | family |
| 11 | 90,045,449 | 90,067,244 | 90,116,848 | 90,122,363 | • 2 COs < 1Mb & NCO > 10kb | 0 | 4 – 11 | family |
| 12 | 50,526,190 | 50,551,529 | 50,564,236 | 50,568,134 | • 2 COs < 1Mb & NCO > 10kb | 1 | 1 – 6 | family |

**Supplementary Table 5**. Positions of the observed crossover events.

| individual | scaffold | position upstream marker | position downstream marker |
|---|---|---|---|
| 1 | 1 | 147,006,260 | 147,798,001 |
| 1 | 1 | 168,779,130 | 171,507,106 |
| 1 | 1 | 177,845,661 | 179,813,558 |
| 1 | 2 | 16,928,257 | 16,964,365 |
| 1 | 2 | 86,438,722 | 86,439,826 |
| 1 | 2 | 129,190,104 | 129,196,955 |
| 1 | 3 | 56,226,446 | 56,228,199 |
| 1 | 3 | 150,842,830 | 150,848,564 |
| 1 | 3 | 174,202,317 | 174,214,869 |
| 1 | 3 | 252,068,274 | 252,070,042 |
| 1 | 4 | 18,189,631 | 18,211,011 |
| 1 | 4 | 76,607,494 | 76,636,477 |
| 1 | 4 | 118,861,702 | 118,881,099 |
| 1 | 4 | 216,521,297 | 216,522,477 |
| 1 | 5 | 13,948,339 | 13,949,447 |
| 1 | 5 | 36,319,680 | 36,323,116 |
| 1 | 5 | 209,746,927 | 209,761,805 |
| 1 | 6 | 10,584,065 | 10,596,898 |
| 1 | 6 | 11,076,447 | 13,501,795 |
| 1 | 6 | 51,785,165 | 51,842,657 |
| 1 | 6 | 57,825,993 | 57,844,742 |
| 1 | 6 | 124,834,502 | 124,861,914 |
| 1 | 6 | 148,174,633 | 148,183,073 |
| 1 | 7 | 2,000,328 | 2,021,107 |
| 1 | 7 | 45,675,558 | 45,737,542 |
| 1 | 7 | 68,666,609 | 68,672,581 |
| 1 | 7 | 80,843,234 | 80,848,607 |
| 1 | 7 | 122,679,946 | 122,721,577 |
| 1 | 7 | 142,762,129 | 142,770,108 |
| 1 | 7 | 185,947,570 | 185,949,405 |
| 1 | 8 | 6,020,034 | 6,031,304 |
| 1 | 8 | 111,071,663 | 111,082,181 |
| 1 | 8 | 153,260,958 | 153,281,425 |
| 1 | 10 | 70,263,131 | 70,277,780 |
| 1 | 10 | 72,567,448 | 72,615,861 |
| 1 | 10 | 85,195,525 | 85,213,693 |
| 1 | 10 | 94,344,717 | 94,346,521 |
| 1 | 10 | 95,167,756 | 95,273,842 |
| 1 | 10 | 101,390,387 | 102,328,886 |
| 1 | 11 | 7,894,999 | 7,899,701 |
| 1 | 11 | 9,510,993 | 9,515,664 |
| 1 | 11 | 80,497,831 | 80,615,600 |
| 1 | 13 | 34,241,677 | 34,250,545 |
| 1 | 13 | 48,444,476 | 48,915,836 |
| 1 | 14 | 31,705,645 | 31,730,633 |
| 2 | 1 | 4,663,484 | 12,503,905 |
| 2 | 1 | 13,314,744 | 13,340,714 |
| 2 | 1 | 22,160,645 | 22,175,120 |
| 2 | 1 | 85,503,373 | 85,519,125 |
| 2 | 1 | 227,355,769 | 227,498,463 |
| 2 | 1 | 285,808,600 | 286,678,903 |
| 2 | 1 | 306,559,801 | 311,093,203 |
| 2 | 2 | 13,452,462 | 13,525,777 |
| 2 | 2 | 133,759,556 | 133,762,027 |
| 2 | 2 | 193,751,972 | 193,760,098 |
| 2 | 2 | 201,341,617 | 202,118,518 |
| 2 | 2 | 281,015,445 | 281,021,771 |
| 2 | 3 | 90,877,579 | 90,886,122 |

| | | | |
|---|---|---|---|
| 2 | 3 | 106,991,952 | 106,994,950 |
| 2 | 3 | 177,566,150 | 179,550,771 |
| 2 | 3 | 179,850,723 | 182,859,186 |
| 2 | 3 | 237,069,090 | 237,080,510 |
| 2 | 3 | 237,564,316 | 245,119,047 |
| 2 | 3 | 255,473,804 | 257,403,995 |
| 2 | 4 | 22,276,215 | 22,286,525 |
| 2 | 4 | 30,945,429 | 30,965,102 |
| 2 | 4 | 111,757,758 | 118,099,593 |
| 2 | 4 | 141,783,444 | 141,786,345 |
| 2 | 4 | 178,237,026 | 178,268,216 |
| 2 | 5 | 13,380,199 | 14,126,225 |
| 2 | 5 | 23,333,811 | 23,336,544 |
| 2 | 5 | 42,811,229 | 42,812,432 |
| 2 | 5 | 127,751,477 | 128,399,868 |
| 2 | 6 | 39,056,246 | 39,060,736 |
| 2 | 6 | 87,307,410 | 87,321,195 |
| 2 | 6 | 109,268,534 | 109,291,683 |
| 2 | 6 | 123,315,217 | 124,083,574 |
| 2 | 6 | 187,205,354 | 187,223,519 |
| 2 | 6 | 192,064,320 | 193,002,569 |
| 2 | 7 | 7,454,396 | 8,252,285 |
| 2 | 7 | 65,907,280 | 67,588,929 |
| 2 | 7 | 83,975,388 | 83,981,923 |
| 2 | 8 | 29,744,149 | 29,783,200 |
| 2 | 8 | 84,872,452 | 84,884,366 |
| 2 | 10 | 3,836,825 | 22,015,129 |
| 2 | 10 | 61,771,729 | 61,772,375 |
| 2 | 10 | 101,837,270 | 101,843,911 |
| 2 | 11 | 29,174,939 | 54,897,338 |
| 2 | 11 | 65,028,461 | 65,031,560 |
| 2 | 11 | 69,027,374 | 69,581,138 |
| 2 | 11 | 82,775,992 | 84,564,074 |
| 2 | 12 | 60,910,242 | 62,442,159 |
| 2 | 12 | 62,734,699 | 65,054,865 |
| 2 | 13 | 19,171,916 | 19,230,498 |
| 2 | 13 | 58,335,038 | 58,356,748 |
| 2 | 14 | 8,715,338 | 8,716,387 |
| 2 | 14 | 21,388,264 | 22,012,546 |
| 2 | 14 | 23,243,395 | 23,269,365 |
| 2 | 15 | 7,565,372 | 7,570,313 |
| 2 | 15 | 9,658,640 | 9,736,730 |
| 3 | 1 | 12,465,768 | 12,504,789 |
| 3 | 1 | 21,204,707 | 21,206,285 |
| 3 | 1 | 89,811,189 | 89,811,645 |
| 3 | 1 | 178,255,559 | 178,264,533 |
| 3 | 1 | 187,405,810 | 188,754,222 |
| 3 | 1 | 218,649,645 | 218,660,485 |
| 3 | 1 | 286,190,572 | 286,222,560 |
| 3 | 1 | 302,823,237 | 302,835,696 |
| 3 | 2 | 7,140,717 | 7,191,100 |
| 3 | 2 | 189,346,065 | 189,350,269 |
| 3 | 2 | 239,602,546 | 239,606,762 |
| 3 | 2 | 284,258,110 | 284,264,321 |
| 3 | 3 | 8,783,507 | 9,000,754 |
| 3 | 3 | 85,968,610 | 85,979,253 |
| 3 | 3 | 175,706,725 | 175,715,035 |
| 3 | 3 | 181,470,348 | 181,494,537 |
| 3 | 3 | 189,298,755 | 189,304,543 |
| 3 | 4 | 16,886,096 | 16,891,573 |
| 3 | 4 | 23,794,131 | 23,798,579 |
| 3 | 4 | 117,283,919 | 117,478,398 |

| | | | |
|---|---|---|---|
| 3 | 4 | 126,196,323 | 126,200,459 |
| 3 | 4 | 165,197,764 | 165,274,408 |
| 3 | 4 | 200,900,130 | 200,944,343 |
| 3 | 4 | 204,161,121 | 204,171,421 |
| 3 | 4 | 215,278,014 | 215,291,723 |
| 3 | 5 | 5,279,766 | 5,778,992 |
| 3 | 5 | 26,511,772 | 26,969,557 |
| 3 | 5 | 31,275,045 | 31,280,660 |
| 3 | 5 | 88,848,062 | 89,068,990 |
| 3 | 6 | 9,261,547 | 9,309,736 |
| 3 | 6 | 117,558,285 | 117,570,845 |
| 3 | 6 | 199,660,926 | 199,697,235 |
| 3 | 7 | 17,325,750 | 17,330,210 |
| 3 | 7 | 22,952,977 | 23,209,325 |
| 3 | 7 | 92,450,549 | 92,480,783 |
| 3 | 7 | 133,261,860 | 133,271,598 |
| 3 | 7 | 164,347,885 | 164,429,072 |
| 3 | 7 | 187,434,698 | 187,437,135 |
| 3 | 8 | 36,260,948 | 36,290,129 |
| 3 | 8 | 152,099,785 | 154,658,132 |
| 3 | 10 | 44,246,996 | 44,252,526 |
| 3 | 10 | 78,849,378 | 79,297,829 |
| 3 | 10 | 89,578,251 | 89,580,021 |
| 3 | 10 | 91,611,006 | 91,612,009 |
| 3 | 11 | 13,389,457 | 13,442,851 |
| 3 | 11 | 50,723,188 | 51,202,493 |
| 3 | 11 | 83,433,913 | 83,606,493 |
| 3 | 11 | 94,008,904 | 95,646,456 |
| 3 | 12 | 6,257,971 | 6,260,918 |
| 3 | 12 | 20,258,993 | 20,262,709 |
| 3 | 13 | 6,626,964 | 6,629,619 |
| 3 | 13 | 46,682,586 | 46,698,633 |
| 3 | 13 | 48,682,435 | 48,683,397 |
| 3 | 13 | 55,845,252 | 55,879,079 |
| 3 | 14 | 7,414,053 | 7,416,655 |
| 3 | 14 | 12,198,476 | 12,343,748 |
| 3 | 14 | 27,063,896 | 27,094,624 |
| 3 | 15 | 22,148,783 | 22,150,157 |
| 4 | 1 | 123,895,618 | 134,078,744 |
| 4 | 1 | 184,920,299 | 184,953,340 |
| 4 | 1 | 286,189,213 | 286,258,513 |
| 4 | 1 | 294,551,566 | 294,565,042 |
| 4 | 1 | 305,397,433 | 305,401,388 |
| 4 | 2 | 26,118,780 | 26,140,130 |
| 4 | 2 | 89,757,671 | 89,770,974 |
| 4 | 2 | 110,958,108 | 113,841,778 |
| 4 | 2 | 173,499,317 | 173,834,558 |
| 4 | 3 | 17,990,991 | 18,963,330 |
| 4 | 3 | 58,438,946 | 58,467,143 |
| 4 | 3 | 104,422,928 | 104,429,151 |
| 4 | 3 | 167,019,951 | 167,489,207 |
| 4 | 3 | 175,216,080 | 175,396,780 |
| 4 | 4 | 35,972,038 | 36,164,529 |
| 4 | 4 | 37,651,245 | 43,809,387 |
| 4 | 4 | 122,608,102 | 122,614,514 |
| 4 | 4 | 145,406,255 | 145,418,682 |
| 4 | 4 | 184,978,678 | 184,992,415 |
| 4 | 4 | 205,756,050 | 205,757,150 |
| 4 | 5 | 27,054,183 | 27,111,879 |
| 4 | 5 | 64,023,013 | 64,178,953 |
| 4 | 5 | 76,594,633 | 76,602,820 |
| 4 | 5 | 108,541,556 | 114,643,431 |

| | | | |
|---|---|---|---|
| 4 | 5 | 147,837,709 | 147,848,249 |
| 4 | 6 | 43,949,661 | 43,958,226 |
| 4 | 6 | 51,482,156 | 55,331,403 |
| 4 | 6 | 101,144,880 | 101,155,340 |
| 4 | 6 | 120,101,662 | 134,620,589 |
| 4 | 6 | 149,863,330 | 152,561,879 |
| 4 | 6 | 173,901,094 | 173,915,402 |
| 4 | 7 | 50,965,271 | 50,967,323 |
| 4 | 7 | 52,515,393 | 52,535,372 |
| 4 | 7 | 167,587,292 | 171,506,176 |
| 4 | 7 | 174,205,751 | 177,079,726 |
| 4 | 8 | 19,497,085 | 19,540,044 |
| 4 | 8 | 37,928,350 | 37,940,200 |
| 4 | 8 | 53,593,282 | 58,098,427 |
| 4 | 8 | 67,701,819 | 67,765,101 |
| 4 | 10 | 52,449,000 | 52,474,199 |
| 4 | 10 | 98,839,912 | 98,893,372 |
| 4 | 10 | 103,699,488 | 103,706,288 |
| 4 | 11 | 25,944,405 | 25,956,207 |
| 4 | 11 | 67,441,697 | 67,446,154 |
| 4 | 11 | 86,160,536 | 86,582,091 |
| 4 | 13 | 3,894,793 | 3,898,951 |
| 4 | 13 | 29,502,261 | 29,507,161 |
| 4 | 13 | 34,248,270 | 34,249,926 |
| 4 | 13 | 39,070,278 | 39,636,357 |
| 4 | 13 | 44,904,176 | 44,923,684 |
| 4 | 13 | 47,586,923 | 47,612,658 |
| 4 | 14 | 17,023,786 | 17,058,876 |
| 4 | 15 | 3,576,660 | 5,569,492 |
| 4 | 15 | 10,643,559 | 10,645,517 |
| 7 | 1 | 183,658,936 | 183,682,006 |
| 7 | 1 | 193,365,344 | 193,389,074 |
| 7 | 1 | 225,881,565 | 225,897,975 |
| 7 | 1 | 282,607,149 | 282,614,263 |
| 7 | 3 | 6,394,039 | 6,396,170 |
| 7 | 3 | 192,871,303 | 192,879,132 |
| 7 | 3 | 243,869,951 | 243,872,771 |
| 7 | 4 | 193,366,845 | 193,380,512 |
| 7 | 5 | 76,606,601 | 76,622,426 |
| 7 | 6 | 61,739,296 | 61,765,894 |
| 7 | 6 | 150,771,636 | 150,772,099 |
| 7 | 6 | 184,703,333 | 186,257,470 |
| 7 | 7 | 8,564,927 | 8,570,476 |
| 7 | 7 | 10,450,043 | 10,490,175 |
| 7 | 7 | 26,481,618 | 26,485,323 |
| 7 | 8 | 67,348,455 | 67,353,609 |
| 7 | 8 | 160,591,602 | 160,592,111 |
| 7 | 11 | 27,649,740 | 27,678,545 |
| 7 | 11 | 95,477,615 | 95,480,298 |
| 7 | 12 | 8,627,169 | 8,643,572 |
| 7 | 13 | 23,587,797 | 23,593,150 |
| 7 | 14 | 11,181,221 | 11,196,349 |
| 7 | 15 | 25,802,419 | 26,058,480 |
| 8 | 1 | 19,717,040 | 19,761,180 |
| 8 | 1 | 54,788,907 | 54,864,578 |
| 8 | 1 | 79,371,272 | 79,383,238 |
| 8 | 1 | 212,312,651 | 212,315,790 |
| 8 | 1 | 220,420,147 | 220,426,338 |
| 8 | 1 | 250,313,174 | 250,315,942 |
| 8 | 1 | 286,192,504 | 286,223,712 |
| 8 | 1 | 301,738,994 | 301,744,149 |
| 8 | 2 | 100,323,136 | 100,362,706 |

| | | | |
|---|---|---|---|
| 8 | 2 | 154,642,231 | 154,649,228 |
| 8 | 2 | 165,023,208 | 165,024,914 |
| 8 | 2 | 268,392,733 | 268,402,746 |
| 8 | 2 | 270,728,566 | 270,743,087 |
| 8 | 3 | 26,045,665 | 26,048,302 |
| 8 | 3 | 28,187,763 | 28,213,753 |
| 8 | 3 | 151,729,232 | 151,743,446 |
| 8 | 3 | 182,559,642 | 182,561,810 |
| 8 | 3 | 183,665,378 | 183,699,778 |
| 8 | 3 | 232,792,960 | 232,793,549 |
| 8 | 4 | 3,564,709 | 3,586,733 |
| 8 | 4 | 147,172,797 | 147,176,182 |
| 8 | 5 | 15,312,725 | 15,343,053 |
| 8 | 5 | 95,218,311 | 95,245,278 |
| 8 | 5 | 190,246,637 | 190,270,685 |
| 8 | 5 | 197,592,716 | 197,650,489 |
| 8 | 5 | 205,704,580 | 205,707,134 |
| 8 | 6 | 13,622,705 | 13,623,071 |
| 8 | 6 | 60,926,287 | 60,948,931 |
| 8 | 6 | 161,050,855 | 161,058,949 |
| 8 | 6 | 197,115,515 | 197,140,320 |
| 8 | 7 | 36,378,597 | 36,391,979 |
| 8 | 7 | 187,701,145 | 187,712,069 |
| 8 | 8 | 63,661,863 | 63,665,742 |
| 8 | 8 | 66,324,631 | 66,344,898 |
| 8 | 8 | 86,775,146 | 86,779,221 |
| 8 | 8 | 109,610,091 | 109,637,459 |
| 8 | 8 | 132,762,980 | 132,781,906 |
| 8 | 10 | 19,827,740 | 19,836,700 |
| 8 | 10 | 52,424,166 | 52,428,778 |
| 8 | 10 | 80,032,319 | 80,033,775 |
| 8 | 10 | 85,199,792 | 85,264,768 |
| 8 | 10 | 90,837,330 | 90,857,615 |
| 8 | 10 | 109,759,368 | 109,774,868 |
| 8 | 11 | 49,045,468 | 49,049,372 |
| 8 | 11 | 76,302,673 | 76,310,116 |
| 8 | 12 | 3,672,385 | 3,757,605 |
| 8 | 12 | 18,448,512 | 18,455,605 |
| 8 | 13 | 27,343,620 | 27,352,628 |
| 8 | 13 | 32,458,356 | 32,471,392 |
| 8 | 13 | 53,972,607 | 53,983,053 |
| 8 | 14 | 10,565,672 | 10,580,778 |
| 8 | 14 | 12,484,605 | 12,486,816 |
| 8 | 15 | 10,662,202 | 10,666,743 |
| 8 | 15 | 16,559,667 | 16,605,290 |
| 9 | 1 | 3,458,801 | 3,469,501 |
| 9 | 1 | 112,525,551 | 112,538,489 |
| 9 | 1 | 220,090,003 | 220,110,407 |
| 9 | 2 | 63,763,496 | 63,771,782 |
| 9 | 2 | 149,013,059 | 149,019,754 |
| 9 | 3 | 65,187,980 | 65,205,156 |
| 9 | 3 | 194,971,740 | 195,021,122 |
| 9 | 4 | 10,501,134 | 10,517,247 |
| 9 | 6 | 149,659,535 | 149,671,043 |
| 9 | 8 | 40,335,411 | 40,343,751 |
| 9 | 8 | 159,024,506 | 159,026,277 |
| 9 | 11 | 37,154,529 | 37,166,409 |
| 9 | 12 | 6,252,762 | 6,271,055 |
| 9 | 12 | 60,660,962 | 60,665,369 |
| 9 | 13 | 47,010,614 | 47,021,563 |
| 9 | 15 | 13,980,653 | 13,987,372 |

**Supplementary Table 6**. Summary statistics of the observed recombination events (*i.e.,* crossovers [COs] and non-crossovers [NCOs]).

| individual | offspring | # CO events | median resolution (bp) | # of NCO events | median tract length (bp) |
|---|---|---|---|---|---|
| 1 | 5 | 14 | 19,932 | 13 | 32 |
| 1 | 6 | 13 | 12,552 | 9 | 1 |
| 1 | 8 | 18 | 12,960 | 11 | 18 |
| 2 | 5 | 19 | 73,315 | 6 | 1 |
| 2 | 6 | 17 | 25,970 | 5 | 1 |
| 2 | 8 | 19 | 18,165 | 9 | 1 |
| 3 | 7 | 11 | 13,709 | 4 | 11 |
| 3 | 9 | 15 | 36,309 | 2 | 1 |
| 3 | 10 | 16 | 11,430 | 5 | 1 |
| 3 | 11 | 16 | 8,427 | 7 | 1 |
| 4 | 7 | 15 | 10,540 | 3 | 1 |
| 4 | 9 | 14 | 256,108 | 1 | 1 |
| 4 | 10 | 12 | 104,700 | 2 | 206 |
| 4 | 11 | 13 | 13,476 | 12 | 1 |
| 7 | 12 | 13 | 13,667 | 16 | 1 |
| 7 | 13 | 10 | 11,121 | 11 | 1 |
| 7 | – | 0 | – | 13 | 1 |
| 8 | 12 | 21 | 9,008 | 18 | 1 |
| 8 | 13 | 15 | 10,013 | 20 | 1 |
| 8 | 14 | 18 | 22,536 | 15 | 1 |
| 8 | – | 0 | – | 4 | 3 |
| 9 | 14 | 16 | 11,229 | 14 | 1 |

**Supplementary Table 7**. Positions and tract lengths of the observed non-crossover events.

| individual | scaffold | first position | last position | tract length (bp) | reference allele | alternate allele |
|---|---|---|---|---|---|---|
| 1 | 1 | 31,716,645 | 31,716,662 | 18 | C,C,G,T,T | A,T,A,C,A |
| 1 | 1 | 83,792,840 | 83,792,840 | 1 | G | C |
| 1 | 1 | 98,931,937 | 98,931,937 | 1 | T | C |
| 1 | 2 | 143,679,725 | 143,679,725 | 1 | C | T |
| 1 | 2 | 173,458,803 | 173,458,934 | 132 | G,C,C,C,C,G,C,T,C,C,T,T,G,G | A,A,T,A,A,T,T,A,T,T,C,G,C,A |
| 1 | 2 | 176,855,468 | 176,856,980 | 1513 | G,T,T,G,G,C,T,T,G,C,G,C,A,G,C,C | A,C,C,A,A,T,C,C,A,A,A,T,C,A,T,A |
| 1 | 2 | 228,634,968 | 228,635,419 | 452 | T,C,T,A,T,T,A,T,T,G,G,T,A,T,T,C,A,<br>G,C,A,T,A,A | A,A,A,G,A,C,G,C,C,T,C,A,C,A,G,T,<br>C,A,T,G,G,G,T |
| 1 | 3 | 52,009,268 | 52,010,136 | 869 | G,C,A,G,T,A,T,A,T,C,C,C,A,T,T,A,G,<br>G,T,C,A,A,C,C,A,T,A | A,T,G,A,A,G,C,G,C,A,T,T,G,A,A,C,<br>A,A,A,T,C,C,A,A,G,C,T |
| 1 | 3 | 229,226,228 | 229,226,240 | 13 | T,T,T | C,C,A |
| 1 | 4 | 25,230,908 | 25,231,189 | 282 | G,T,G,G,G,G,G,C,G,C,G,C,C,T,A,G,<br>G,C,T,G,T,G,G,G,G | A,A,A,T,A,C,A,T,A,A,A,T,A,C,G,A,<br>C,A,C,A,C,A,A,A,A |
| 1 | 4 | 108,863,434 | 108,863,434 | 1 | C | T |
| 1 | 4 | 114,580,815 | 114,581,127 | 313 | T,C,C,G,G,G,G,C,C,G,G,C,C,G,A,G,<br>T,C,G | G,T,A,A,A,T,T,G,T,C,T,T,T,C,G,A,<br>C,T,C |
| 1 | 5 | 15,591,979 | 15,592,357 | 379 | G,A,C | A,G,T |
| 1 | 6 | 24,809,637 | 24,809,709 | 73 | T,T,C,C,G,G,T,T,T | C,A,T,T,T,A,C,G,C |
| 1 | 6 | 51,475,887 | 51,475,887 | 1 | C | T |
| 1 | 6 | 175,795,974 | 175,796,638 | 665 | T,T,A,A,T,T,G,T,T,C,C | C,C,T,G,C,C,A,C,A,T,A |
| 1 | 7 | 5,832,693 | 5,832,693 | 1 | C | G |
| 1 | 7 | 11,889,983 | 11,890,022 | 40 | A,C,T | C,T,C |
| 1 | 7 | 137,839,910 | 137,841,044 | 1135 | A,T,T,G,C,C,C,T,C,C,C,A,A,G,G,G,<br>C,T,T,C,C,C,T,G,T,T,G,T,T,C,C,C,G,<br>G,A,C,G,A,C,A,A,A,T,G,A,C,C,A,C,A,<br>G,G,A,G,G,A,A,T | G,C,C,A,A,T,T,C,T,T,T,G,G,A,A,A,<br>T,C,C,G,G,T,C,A,C,C,A,C,G,A,T,G,<br>T,A,G,A,C,G,T,G,T,G,A,T,G,T,T,G,<br>T,G,T,A,G,A,A,C,T,C |
| 1 | 7 | 164,612,573 | 164,612,573 | 1 | C | G |
| 1 | 7 | 166,146,917 | 166,146,917 | 1 | C | T |
| 1 | 8 | 158,263,527 | 158,263,527 | 1 | G | A |
| 1 | 10 | 34,607,984 | 34,607,984 | 1 | G | A |
| 1 | 10 | 97,749,510 | 97,749,541 | 32 | C,A,C,T,A | A,G,T,G,G |
| 1 | 11 | 5,629,608 | 5,630,187 | 580 | C,A,C,C,A,G,T,G,T,G,G,A,T,C,A,G,<br>G,G,T,C,G,G,G | A,T,G,T,G,A,C,A,A,A,C,G,C,T,T,C,<br>A,A,C,T,T,A,A |
| 1 | 11 | 78,417,901 | 78,417,933 | 33 | C,G,C,G,T,C | T,A,G,A,C,T |
| 1 | 11 | 78,417,987 | 78,417,987 | 1 | C | T |
| 1 | 12 | 8,295,111 | 8,295,127 | 17 | C,G,T | T,A,A |
| 1 | 12 | 24,970,285 | 24,970,285 | 1 | G | A |
| 1 | 12 | 38,286,016 | 38,286,016 | 1 | T | C |
| 1 | 13 | 47,189,693 | 47,189,699 | 7 | C,C | T,T |
| 1 | 14 | 12,592,451 | 12,592,451 | 1 | G | A |
| 1 | 15 | 13,247,585 | 13,247,813 | 229 | C,G,T,T,G,T,G,T,G,T,A,C,G | T,T,G,C,A,A,A,C,A,C,T,T,T |
| 2 | 1 | 261,273,346 | 261,273,346 | 1 | A | G |

| | | | | | | |
|---|---|---|---|---|---|---|
| 2 | 1 | 281,720,785 | 281,724,890 | 4106 | G,C,G | A,T,T |
| 2 | 1 | 303,213,916 | 303,213,916 | 1 | A | T |
| 2 | 2 | 76,418,473 | 76,418,473 | 1 | G | C |
| 2 | 3 | 175,013,127 | 175,013,127 | 1 | C | G |
| 2 | 3 | 175,288,959 | 175,288,959 | 1 | A | C |
| 2 | 4 | 10,276,389 | 10,276,389 | 1 | A | G |
| 2 | 4 | 193,029,181 | 193,032,980 | 3800 | G,T,G,T | A,A,A,C |
| 2 | 7 | 8,310,569 | 8,310,569 | 1 | T | C |
| 2 | 7 | 10,207,374 | 10,207,374 | 1 | C | T |
| 2 | 11 | 20,836,749 | 20,836,749 | 1 | C | T |
| 2 | 12 | 16,767,242 | 16,767,242 | 1 | A | C |
| 2 | 12 | 16,780,464 | 16,780,464 | 1 | G | A |
| 2 | 12 | 17,342,535 | 17,342,535 | 1 | A | G |
| 2 | 12 | 17,349,923 | 17,349,923 | 1 | A | C |
| 2 | 12 | 19,090,200 | 19,090,200 | 1 | C | T |
| 2 | 13 | 49,073,484 | 49,073,527 | 44 | G,T,T,T,C,A | C,C,C,C,T,C |
| 2 | 13 | 51,852,078 | 51,852,078 | 1 | A | G |
| 2 | 14 | 11,338,111 | 11,338,111 | 1 | T | G |
| 2 | 14 | 20,817,007 | 20,817,007 | 1 | A | C |
| 3 | 1 | 97,125,790 | 97,125,790 | 1 | T | C |
| 3 | 1 | 135,122,088 | 135,122,088 | 1 | G | A |
| 3 | 1 | 231,896,237 | 231,896,237 | 1 | T | C |
| 3 | 1 | 303,937,881 | 303,937,881 | 1 | T | C |
| 3 | 2 | 261,997,843 | 261,997,843 | 1 | C | A |
| 3 | 3 | 44,012,071 | 44,012,071 | 1 | G | A |
| 3 | 3 | 96,810,249 | 96,810,262 | 14 | G,C,A | A,T,G |
| 3 | 3 | 163,757,163 | 163,757,163 | 1 | G | A |
| 3 | 3 | 170,285,829 | 170,285,829 | 1 | G | A |
| 3 | 4 | 72,852,544 | 72,852,544 | 1 | A | C |
| 3 | 4 | 114,580,931 | 114,581,429 | 499 | G,C,C,G,A,G,T,C,G,C,C,C,T,G,C,C,A,C,A,G,T | T,T,T,C,G,A,C,T,C,G,T,T,C,T,T,T,G,T,C,A,A |
| 3 | 5 | 26,071,801 | 26,072,976 | 1176 | G,A,T,C,A,G,G,A,C,T,A,A,T,T,C,A,A,G,T | T,C,C,G,G,A,A,G,T,G,T,T,C,G,T,G,G,A,C |
| 3 | 5 | 150,510,916 | 150,510,966 | 51 | G,G,T,A,C | A,A,C,T,T |
| 3 | 5 | 191,979,105 | 191,979,105 | 1 | A | G |
| 3 | 7 | 93,987,902 | 93,987,902 | 1 | T | A |
| 3 | 10 | 86,908,867 | 86,908,867 | 1 | A | G |
| 3 | 11 | 5,619,725 | 5,619,744 | 20 | C,G | T,A |
| 3 | 12 | 17,234,966 | 17,234,966 | 1 | A | T |
| 4 | 1 | 19,720,998 | 19,720,998 | 1 | G | A |
| 4 | 1 | 104,181,989 | 104,181,989 | 1 | T | G |
| 4 | 3 | 76,628,762 | 76,628,762 | 1 | A | G |
| 4 | 4 | 2,252,960 | 2,253,493 | 534 | G,G,C,A,C | C,T,A,G,T |
| 4 | 7 | 148,255,538 | 148,255,538 | 1 | C | A |
| 4 | 7 | 190,148,575 | 190,148,575 | 1 | T | C |

| | | | | | | |
|---|---|---|---|---|---|---|
| 4 | 7 | 197,026,266 | 197,026,266 | 1 | G | A |
| 4 | 8 | 76,288,035 | 76,288,035 | 1 | A | T |
| 4 | 8 | 77,694,955 | 77,694,960 | 6 | C,G | A,A |
| 4 | 8 | 82,879,163 | 82,879,163 | 1 | G | A |
| 4 | 8 | 152,863,076 | 152,863,090 | 15 | G,C,A | A,T,G |
| 4 | 10 | 80,517,412 | 80,517,821 | 410 | G,A | A,G |
| 4 | 12 | 2,284,211 | 2,284,211 | 1 | A | G |
| 4 | 12 | 50,564,235 | 50,566,491 | 2257 | A,C,A,C | G,A,G,T |
| 4 | 12 | 62,443,480 | 62,443,480 | 1 | A | G |
| 4 | 13 | 48,068,797 | 48,068,797 | 1 | A | G |
| 4 | 15 | 6,067,222 | 6,067,222 | 1 | G | C |
| 4 | 15 | 20,863,479 | 20,863,479 | 1 | A | G |
| 7 | 1 | 98,931,872 | 98,931,960 | 89 | G,C,T,A | A,G,C,G |
| 7 | 1 | 121,350,416 | 121,350,427 | 12 | G,C,G | T,T,C |
| 7 | 1 | 125,057,576 | 125,057,608 | 33 | C,A | T,G |
| 7 | 1 | 146,266,013 | 146,266,037 | 25 | T,A,T,T | C,G,C,C |
| 7 | 1 | 146,351,177 | 146,351,201 | 25 | C,G,G,C | T,A,A,G |
| 7 | 1 | 146,516,813 | 146,516,813 | 1 | C | T |
| 7 | 1 | 146,563,366 | 146,563,366 | 1 | G | T |
| 7 | 1 | 209,783,906 | 209,783,906 | 1 | G | A |
| 7 | 1 | 262,631,373 | 262,631,373 | 1 | C | T |
| 7 | 1 | 262,763,300 | 262,763,300 | 1 | A | G |
| 7 | 1 | 313,019,108 | 313,019,108 | 1 | C | T |
| 7 | 2 | 156,820,917 | 156,821,256 | 340 | G,G,T,G,C,T,C,C,T,C,T,T,A,G,C,C,A | T,C,A,A,G,A,T,T,G,T,A,A,G,A,T,T,G |
| 7 | 2 | 173,458,593 | 173,459,012 | 420 | C,G,C,G,C,C,C,C,G,C,T,C,C,T,T,G,G,C,T,C,T,G,A | A,C,A,A,A,T,A,A,T,T,A,T,T,C,G,C,A,T,C,T,C,A,G |
| 7 | 2 | 176,855,492 | 176,857,011 | 1520 | T,G,G,C,T,T,G,C,G,C,A,G,C,C,T,T,T | C,A,A,T,C,C,A,A,A,T,C,A,T,A,A,C,C |
| 7 | 2 | 228,634,889 | 228,635,697 | 809 | C,G,A,C,A,C,G,T,C,T,A,T,T,A,T,T,G,G,T,A,T,T,C,A,G,C,A,T,A,A,T | T,T,T,G,G,T,A,A,A,A,G,A,C,G,C,C,T,C,A,C,A,G,T,C,A,T,G,G,G,T,C |
| 7 | 3 | 19,059,687 | 19,059,687 | 1 | G | A |
| 7 | 3 | 52,008,409 | 52,010,157 | 1749 | A,G,C,G,T,A,T,C,C,G,A,A,G,G,G,C,A,G,T,A,T,A,T,C,C,C,A,T,T,A,G,G,T,C,A,A,C,C,A,T,A,T,C | C,A,T,A,C,G,A,T,T,A,C,G,C,A,A,T,G,A,A,G,C,G,C,A,T,T,G,A,A,C,A,A,A,T,C,C,A,A,G,C,T,C,A |
| 7 | 3 | 65,155,382 | 65,155,385 | 4 | T,A | C,G |
| 7 | 3 | 150,047,395 | 150,047,395 | 1 | G | A |
| 7 | 3 | 170,068,701 | 170,068,839 | 139 | G,G,C,T,A | A,C,T,C,G |
| 7 | 4 | 4,018,076 | 4,018,076 | 1 | A | C |
| 7 | 4 | 23,016,152 | 23,016,152 | 1 | G | A |
| 7 | 4 | 180,976,963 | 180,976,963 | 1 | T | C |
| 7 | 5 | 62,518,736 | 62,518,767 | 32 | C,T,C,A,C | G,C,T,G,A |
| 7 | 5 | 142,085,964 | 142,085,964 | 1 | T | C |
| 7 | 5 | 156,424,845 | 156,424,845 | 1 | G | A |
| 7 | 5 | 194,075,710 | 194,075,710 | 1 | G | A |

| | | | | | | |
|---|---|---|---|---|---|---|
| 7 | 5 | 200,142,207 | 200,142,207 | 1 | G | T |
| 7 | 7 | 6,182,210 | 6,182,210 | 1 | G | A |
| 7 | 7 | 120,370,136 | 120,370,136 | 1 | C | A |
| 7 | 7 | 121,210,670 | 121,210,670 | 1 | C | A |
| 7 | 7 | 154,164,950 | 154,164,950 | 1 | C | G |
| 7 | 8 | 11,294,318 | 11,294,318 | 1 | T | A |
| 7 | 8 | 102,549,277 | 102,549,277 | 1 | G | A |
| 7 | 11 | 7,598,732 | 7,598,732 | 1 | C | T |
| 7 | 11 | 9,644,910 | 9,644,910 | 1 | T | G |
| 7 | 11 | 68,500,035 | 68,502,218 | 2184 | C,T,A,C,T,G,G,T,C,A,T,A,C,G,G,G,T, A,C,T,C,G,T,T,C,C,A,A,A,T,C,C,G,C, G,C,G,G,G,C,C,A,G,G,G,C,C,T,C,G, G,G,C,C,C,C,G,G,C,G,T,G,G,A,A,C, C,C,T,G,T,A,A,C | A,G,G,A,G,C,A,C,A,G,A,G,T,A,A,C, C,T,A,A,A,C,A,C,A,T,C,T,G,C,T,T, A,A,T,T,C,T,T,A,T,G,A,A,T,A,T,A,A, A,A,A,A,A,A,A,A,A,T,A,C,A,A,T,G, T,T,T,C,A,C,C,T,T |
| 7 | 12 | 8,294,976 | 8,295,330 | 355 | G,A,T,G,G,T,C,G,T,C,T,G,G,G,T | C,T,C,A,A,A,T,A,A,T,C,A,T,A,G |
| 7 | 13 | 51,849,268 | 51,849,268 | 1 | A | G |
| 7 | 14 | 29,122,668 | 29,122,668 | 1 | A | G |
| 8 | 1 | 117,155,935 | 117,155,935 | 1 | T | C |
| 8 | 1 | 133,389,710 | 133,389,951 | 242 | T,C,C | A,T,T |
| 8 | 1 | 194,960,564 | 194,960,564 | 1 | C | T |
| 8 | 1 | 208,283,335 | 208,283,335 | 1 | C | G |
| 8 | 1 | 231,950,492 | 231,950,492 | 1 | G | T |
| 8 | 1 | 269,983,738 | 269,983,738 | 1 | C | T |
| 8 | 2 | 103,775,996 | 103,775,996 | 1 | T | C |
| 8 | 2 | 139,349,418 | 139,349,426 | 9 | G,C,A | A,T,C |
| 8 | 2 | 139,511,269 | 139,511,269 | 1 | C | T |
| 8 | 2 | 139,523,820 | 139,523,820 | 1 | G | A |
| 8 | 2 | 185,361,856 | 185,361,856 | 1 | C | T |
| 8 | 3 | 96,765,469 | 96,765,469 | 1 | C | T |
| 8 | 3 | 113,132,887 | 113,132,887 | 1 | A | G |
| 8 | 3 | 168,089,964 | 168,089,964 | 1 | A | G |
| 8 | 3 | 178,147,003 | 178,147,003 | 1 | A | T |
| 8 | 3 | 232,338,634 | 232,338,634 | 1 | A | G |
| 8 | 4 | 119,355,646 | 119,355,646 | 1 | A | G |
| 8 | 4 | 127,619,912 | 127,619,912 | 1 | A | C |
| 8 | 4 | 136,930,605 | 136,930,605 | 1 | C | G |
| 8 | 4 | 171,383,322 | 171,383,322 | 1 | G | A |
| 8 | 5 | 72,696,781 | 72,696,781 | 1 | T | C |
| 8 | 5 | 81,570,384 | 81,570,384 | 1 | G | A |
| 8 | 5 | 130,005,093 | 130,005,093 | 1 | C | T |
| 8 | 5 | 142,705,414 | 142,705,414 | 1 | G | A |
| 8 | 5 | 197,641,066 | 197,641,066 | 1 | C | T |
| 8 | 6 | 22,785,435 | 22,785,435 | 1 | T | C |
| 8 | 6 | 32,299,798 | 32,299,798 | 1 | C | T |
| 8 | 6 | 169,321,580 | 169,321,580 | 1 | A | G |

| | | | | | | |
|---|---|---|---|---|---|---|
| 8 | 6 | 196,201,741 | 196,201,741 | 1 | C | T |
| 8 | 6 | 200,196,214 | 200,196,214 | 1 | G | A |
| 8 | 6 | 201,237,027 | 201,237,758 | 732 | G,A | A,C |
| 8 | 7 | 85,274,442 | 85,274,442 | 1 | A | C |
| 8 | 7 | 88,369,946 | 88,369,946 | 1 | C | G |
| 8 | 7 | 118,656,911 | 118,656,911 | 1 | C | T |
| 8 | 7 | 165,906,822 | 165,906,822 | 1 | G | A |
| 8 | 8 | 53,239,233 | 53,239,233 | 1 | C | T |
| 8 | 8 | 57,953,435 | 57,953,435 | 1 | G | C |
| 8 | 8 | 58,098,599 | 58,098,599 | 1 | G | A |
| 8 | 8 | 70,609,820 | 70,609,820 | 1 | C | T |
| 8 | 10 | 109,983,094 | 109,983,094 | 1 | A | C |
| 8 | 11 | 22,823,112 | 22,823,112 | 1 | A | G |
| 8 | 11 | 41,012,162 | 41,012,162 | 1 | G | A |
| 8 | 11 | 69,463,844 | 69,463,844 | 1 | T | A |
| 8 | 11 | 82,379,647 | 82,379,667 | 21 | C,G,G,A,A | G,A,C,T,G |
| 8 | 11 | 83,643,914 | 83,643,914 | 1 | G | T |
| 8 | 12 | 3,005,022 | 3,005,022 | 1 | T | G |
| 8 | 12 | 15,208,040 | 15,208,040 | 1 | C | T |
| 8 | 12 | 16,295,847 | 16,295,847 | 1 | C | G |
| 8 | 12 | 22,253,507 | 22,253,507 | 1 | T | C |
| 8 | 12 | 31,615,778 | 31,615,778 | 1 | C | T |
| 8 | 12 | 42,418,606 | 42,418,606 | 1 | T | G |
| 8 | 12 | 62,359,774 | 62,359,774 | 1 | T | G |
| 8 | 13 | 3,044,597 | 3,046,373 | 1777 | G,C,C,C,C,C,C,C | A,T,T,T,T,G,T,T |
| 8 | 13 | 14,373,063 | 14,373,063 | 1 | T | C |
| 8 | 13 | 47,188,974 | 47,189,816 | 843 | A,C,A,T,G,C,G,T,C,A,A,A,G,G,A,G, A,T,C,C,T,A,G,G,G | C,A,C,C,T,G,C,C,T,G,G,G,A,T,T,C, C,C,T,T,A,C,A,C,A |
| 8 | 13 | 49,298,407 | 49,298,407 | 1 | C | T |
| 8 | 13 | 51,839,564 | 51,839,568 | 5 | T,G,T | C,A,C |
| 9 | 1 | 186,465,304 | 186,465,304 | 1 | G | T |
| 9 | 1 | 226,013,212 | 226,013,231 | 20 | G,A,C | A,G,T |
| 9 | 2 | 173,458,865 | 173,458,890 | 26 | C,C,C,G | T,A,A,T |
| 9 | 2 | 226,139,860 | 226,139,943 | 84 | G,T,C,C,A,A,T,C,G | C,C,T,G,G,T,C,T,C |
| 9 | 3 | 115,269,157 | 115,269,157 | 1 | G | T |
| 9 | 3 | 132,829,084 | 132,829,084 | 1 | G | A |
| 9 | 3 | 176,898,998 | 176,898,998 | 1 | C | T |
| 9 | 5 | 150,490,837 | 150,490,949 | 113 | T,C,A,T,C,T,C,C,T,G,G,G,G | G,T,T,C,G,C,T,T,C,C,A,C,C |
| 9 | 5 | 212,152,293 | 212,152,293 | 1 | T | A |
| 9 | 6 | 74,457,871 | 74,457,871 | 1 | A | G |
| 9 | 6 | 175,795,872 | 175,796,518 | 647 | A,T,G,T,C,G,T,C,T,T,A,A,T,T | G,G,A,G,T,T,C,T,C,C,T,G,C,C |
| 9 | 8 | 98,646,538 | 98,646,607 | 70 | G,G,C,T,C | A,C,T,A,T |
| 9 | 10 | 38,869,502 | 38,869,502 | 1 | C | T |
| 9 | 12 | 62,638,517 | 62,638,517 | 1 | T | C |

**Supplementary Figure 1**.

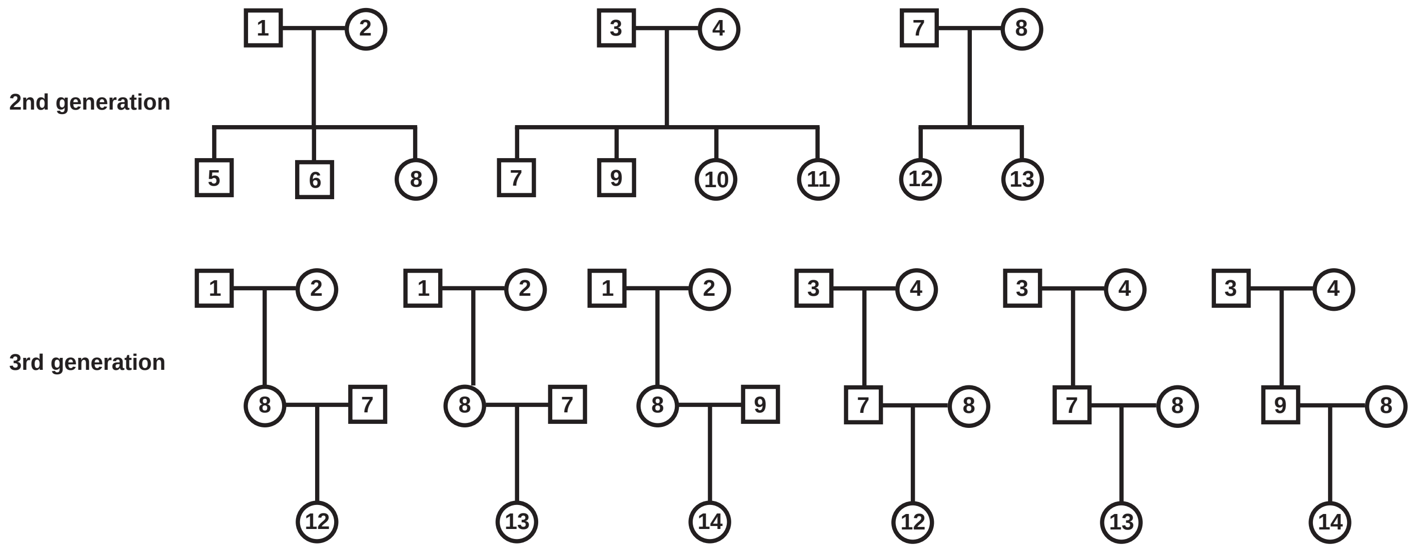

Structure of the three two-generation nuclear families with multiple offspring (top) and six three-generation pedigrees (bottom) in which gamete transmission was tracked to identify recombination events in aye-ayes.

**Supplementary Figure 2**.

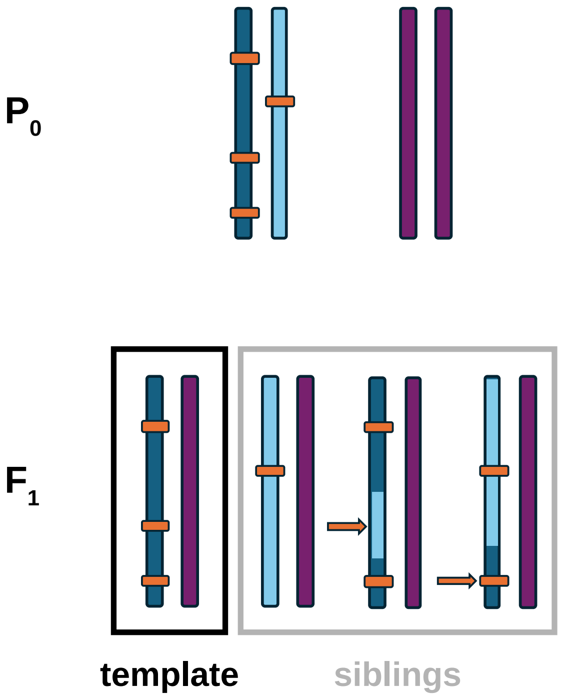

A schematic representation of the recombination detection approach in a two-generation family. Highlighted by orange boxes are phase-informative markers that can be used to detect recombination events inherited by the $F_1$ individuals by comparing the haplotype of the randomly assigned "template" offspring to each of its siblings.

**Supplementary Figure 3**.

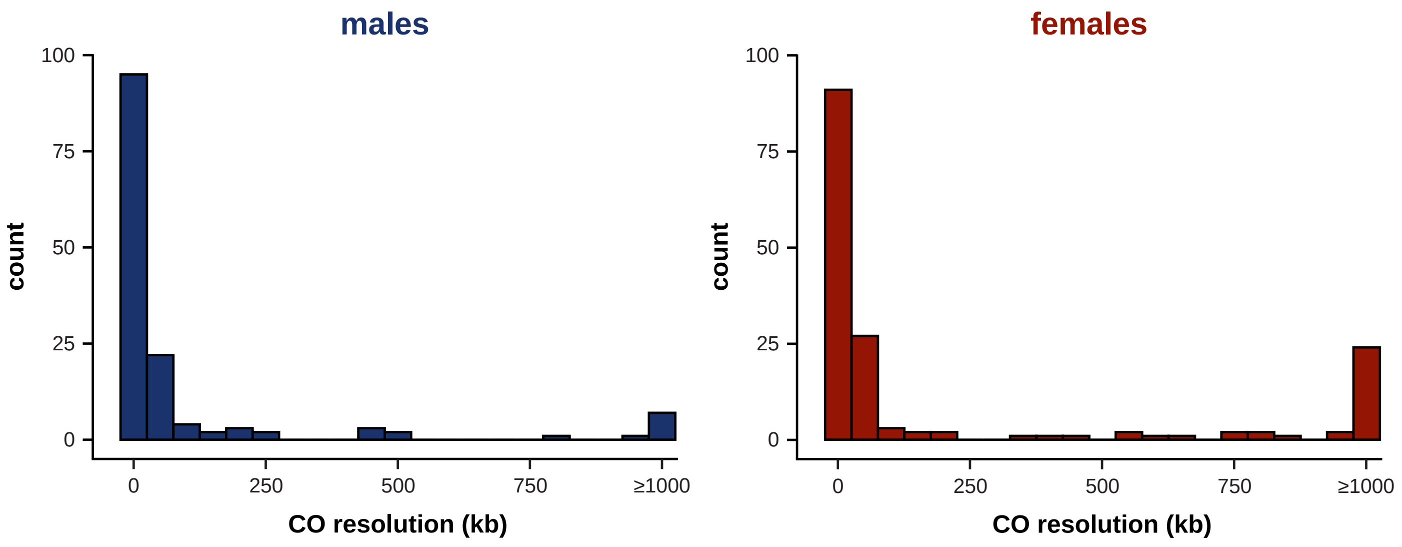

Distribution of crossover (CO) resolution (reported on a kilobase [kb]-scale) based on the two closest informative markers flanking each event in the 10 maternal and 10 paternal meioses.

**Supplementary Figure 4**.

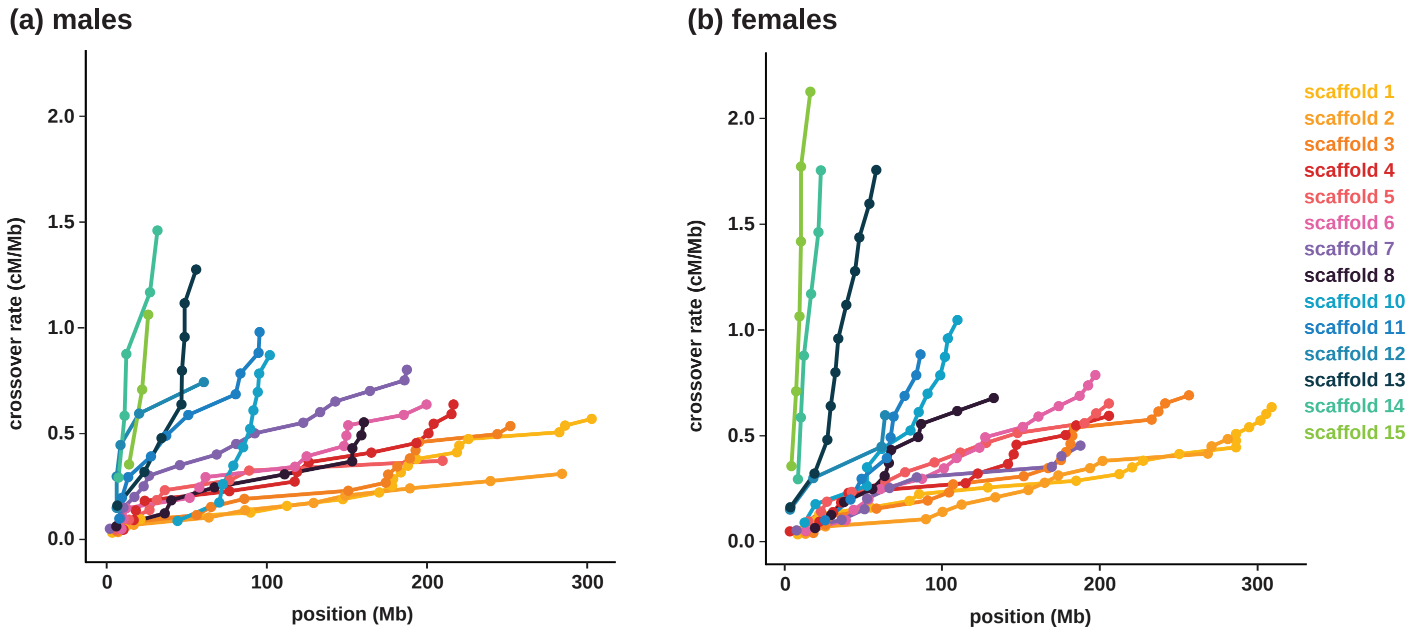

Relationship between the cumulative genetic distance obtained from the distribution of crossover events and the physical length of each autosome in (a) males and (b) females (scaffold 9 is not shown as it corresponds to the X-chromosome).

**Supplementary Figure 5**.

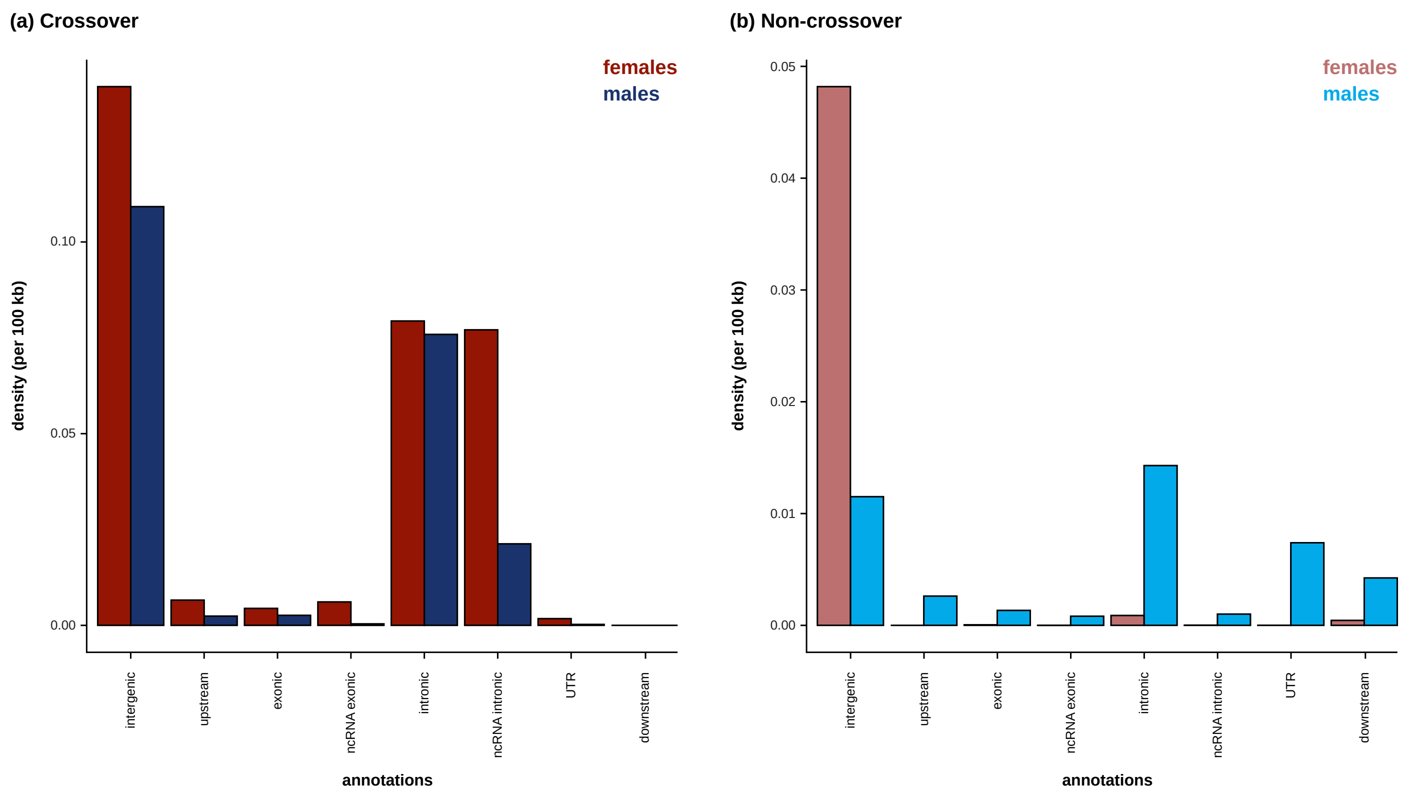

Density of sex-specific (a) crossover events with a resolution of less than 5 kb and (b) non-crossover events by genomic region (*i.e.*, intergenic, upstream, exonic, exonic non-coding RNA [ncRNA], intronic, intronic ncRNA, 3' and 5' UTR, and downstream).
